# Glial cells gate small RNA transport from germline to brain during transgenerational epigenetic inheritance of behavior

**DOI:** 10.64898/2026.10.24.761599

**Authors:** Abigail K. Brown, Xinbei Li, Rachel Kaletsky, Renee Seto, Katherine Novak, Liyang Sun, Coleen T. Murphy

## Abstract

How circulating RNAs reach their target tissue is not well understood. In *C. elegans*, small RNA signals that convey exposure to pathogenic *Pseudomonas* travel from the intestine to germline to neurons, allowing both learned pathogen avoidance and the transgenerational inheritance of this information for four generations – but how this Cer1 retrotransposon-bound RNA signal reaches the nervous system was previously unknown. We found that the RNA transporter SID-1, long thought to act in the intestine or germline, instead functions in a single cell type, the amphid sheath glia, where it is necessary and sufficient for learned avoidance. SID-1 mediates uptake of germline-derived RNA, followed by transcytosis to a neighboring sensory neuron, where the P11 target gene *maco-1* is silenced. This glial “filter”, which uses highly conserved molecular components to transcytose RNAs to neurons, reveals a general mechanism for gating small RNA access to the nervous system, paralleling the mammalian blood-brain barrier.

## INTRODUCTION

All animals need to interpret signals from their environment to distinguish positive cues, such as food and mates, from negative stimuli, including pathogens. *C. elegans* demonstrates a curious behavior: they are first attracted to the bacterial pathogen *Pseudomonas aeruginosa* (PA14) before learning to avoid it^1–3^, likely because many *Pseudomonas* species, both food sources and pathogens, smell similarly to *C. elegans*^4^. We previously found that worms can learn to avoid several pathogenic *Pseudomonas* species^3–5^ by interpreting specific small RNA signals made by the bacteria^6^. These small RNAs have 16- or 17nt perfect sequence matches the neuronal genes *maco-1* and *vab-1*, which are subsequently downregulated, suggesting an underlying RNA interference-like mechanism^4–6^; this model is supported by the requirement for RNA transporters, RNA-dependent polymerase, and Argonautes for this RNA-mediated learned avoidance^3,6,7^. Remarkably, ingestion and processing of these bacterial small RNAs results in the inheritance of learned pathogen avoidance for four generations^3,6,4,5^. We also discovered that virus-like particles of the Ty3/Gypsy retrotransposon Cer1 are required for this phenomenon, acting at the step of communication from germline to the nervous system^8^; these RNA-carrying Cer1 VLPs can also transmit learned information horizontally from trained to naive animals^8^.

Here, we focus on the P11-*maco-1* arm of this pathway to dissect the cellular route of RNA transmission. Because of parallels with RNA interference, we previously tested mutants of the genes in the RNAi pathway, including the dsRNA transporter SID-1, the transmembrane protein SID-2, and the endoribonuclease DCR-1/Dicer, and found that loss of each of these genes abolished small RNA-induced learned pathogen avoidance^4–6^. SID-2 takes up environmental RNAs into the intestine^9,10^, where Dicer cleaves the RNA^11^; we confirmed through rescue experiments that Dicer functions in the intestine in the P11 pathway^6^. Available literature implied that SID-1 likely acts to transport sRNA fragments from the intestine into the germline^9,12,13^. However, we noted that unlike *sid-2*, *dcr-1,* and germline mutants (e.g., *prg-1, rrf-1, rrf-3* and *meg*), which lack the ability to learn avoidance when trained on P11 sRNA but have normal naive attraction to PA14, *sid-1* mutants exhibit high naive avoidance of PA14 in addition to lacking P11-learned avoidance – a phenotype also exhibited by *maco-1* mutants^6^ and Cer1 mutants^8^. That is, *sid-1* and *maco-1* mutants both avoid PA14 without prior exposure to the pathogen or its small RNA, unlike mutants of genes that normally function in the intestine or germline.

*sid-1* (systemic RNA interference defective) was first identified for its role in systemic RNAi^13^, and a model for its function was established: SID-1 was proposed to function as a channel enabling passive, non-endocytic uptake of large dsRNA specifically in non-neuronal tissues, primarily the intestine and germline^13–15^. However, recent cryo-EM structures and biochemical analyses of SID-1 and its human orthologs SIDT1 and SIDT2 have indicated important revisions to this model, including that RNA uptake occurs via an endocytic rather than channel-based mechanism and that RNA substrates considerably shorter than originally proposed can be bound by SID-1^16–23^. Whether the tissue-restriction component of this model also requires revision - and, if so, where SID-1 actually functions, whether it requires endocytosis or not, and whether it acts in an RNA-dependent manner - has not been tested.

Here, using bacterial small RNA-induced learned pathogen avoidance as our paradigm, we find that SID-1 is dispensable in the germline and intestine, and instead functions solely in a single cell type of the nervous system, the amphid sheath glia (AMsh), where it is both necessary and sufficient for this behavior. Our findings suggest a new model for SID-1 function, one in which amphid sheath glial SID-1 plays a critical role in the interpretation of environmental RNA information.

## RESULTS

### sid-1 is required for sRNA-induced learned pathogen avoidance

We previously found that *C. elegans* trained either with *Pseudomonas aeruginosa* PA14 or its small RNA, P11, learn to avoid the pathogen, and this learned avoidance is transmitted to four generations of its progeny^3,6^. We also found that this effect required the activity of SID-1, the systemic RNAi transporter ^4–6^; in fact, *sid-1* mutants show high naive avoidance of the pathogen, and cannot learn to avoid PA14 further upon P11 training (Fig. 1A, B). Furthermore, *sid-1* mutants do not exhibit the increase in short reversals that wild-type worms display after P11 sRNA training^5^ (Fig. 1C), or the increase in expression of *daf-7p::gfp* in the ASI that normally follows P11 training (Fig. 1D, E). Together, these data show that SID-1 is a necessary component of the naive setpoint of PA14 avoidance and is required for other P11-induced phenotypes.

**Figure 1:**
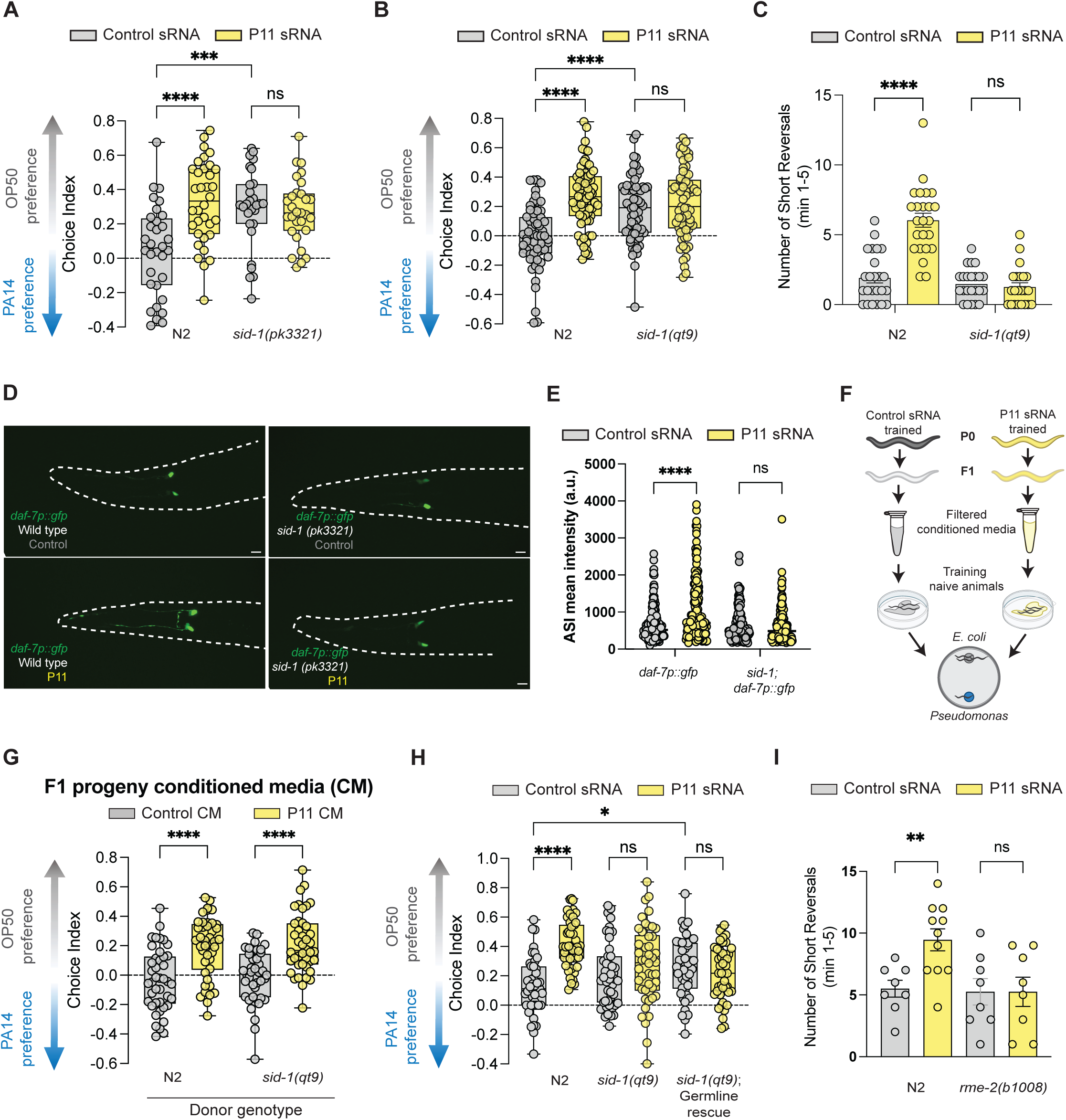
*sid-1* is required for proper P11-induced learned avoidance, and does not function in germline, intestine, or neurons. A, B. Wild-type (N2) and *sid-1* mutant animals were treated with P11 or control sRNA bacteria for 24 hours. *sid-1(pk3321)* (A) and *sid-1(qt9)* (B) exhibit high naive avoidance of PA14 and cannot learn to avoid PA14 with P11 sRNA training. Choice index = (# animals on OP50 - # animals on PA14)/ (total # of animals). C. The number of short reversals is increased with P11 sRNA training in wild type animals and was not observed with P11 sRNA training in *sid-1* animals. Three independent biological replicates were quantified. D. *sid-1* mutants do not exhibit the increase in expression of *daf-7p::gfp* in the ASI that accompanies P11 training. Representative images of 24-hour P11 or control bacteria trained wild-type and sid-1 worms expressing *daf-7p::gfp* in the ASI neurons. Scale bars are 100μm. E. Scatter plot quantification of daf-7p::gfp in ASI neurons show wild type and sid-1 worms following 24-hour exposure to control and P11 sRNA. Three independent biological replicates were quantified. Two-way ANOVA with Tukey’s multiple comparisons test. \*\*\*\**p<*0.0001, ns *p>*0.9999 not significant. F. Schematic of the horizontal transfer assay. Training naive animals with conditioned media from F1 progeny wild type and *sid-1* animals trained with control or P11 sRNA. G. Media of F1s from *sid-1* mutant worms trained on P11 sRNA had no defect in ability to horizontally transfer learned information. H. Expression of *sid-1* in the germline rescue does not rescue learned pathogen avoidance. I. *rme-2* mutants are unable to increase short reversals upon P11 training. For choice assays each dot represents an assay plate of 20-194 worms, average = 54 worms /plate, average of 15 plates/genotype/training condition; n ≥ 3 biological replicates. Boxplots: center line represents median; box range denotes the 25th to 75th percentile; whiskers represent minimum to maximum. Two-way ANOVA with Tukey’s multiple comparison’s test (A-B, E-F, I, L). \*\*\*\**p<*0.0001, \*\*\**p<*0.001, \*\**p<*0.01, \**p<*0.05, ns = not significant. Scatter plot: each dot represents a neuron; center line denotes median.

### SID-1 function is not required in the germline or intestine for sRNA-induced learned pathogen avoidance

Previously published data suggested that SID-1 functions in the germline^13^. To test this hypothesis, we performed an experiment that distinguishes genes required upstream of Cer1 generation of cargo-packed VLPs from those that act downstream to execute behavior^5,8^: we treated naive worms with media conditioned with the F1s of trained animals (Fig. 1F); conditioned media of mutants in genes acting in the initiation (intestine) or propagation (germline) steps of transgenerational inheritance do not confer learned avoidance upon naive animals^8^. However, we observed that media conditioned with F1s from *sid-1* mutant worms trained on P11 sRNA had no defect in the ability to horizontally transfer learned information (Fig. 1G), suggesting that SID-1 does not act in the intestine or germline. To further test this model, we rescued *sid-1* expression in the germline of the *sid-1(qt9)* mutant, but we found that this did not rescue P11 learned avoidance behavior or naive attraction (Fig. 1H). Together, these data rule out a role for SID-1 in either the intestine or the germline in P11-mediated learned avoidance. These results instead suggest that SID-1 must be required in a tissue downstream of the germline.

If SID-1 is not required for RNA entry from the intestine, another protein must carry out this role. The yolk/vitellogenin low-density lipoprotein receptor RME-2 has been previously shown to facilitate RNA entry into the germline^24,25^. Because *rme-2* mutants are severely sick and cannot be reliably obtained in the numbers required for population-level choice assays (typically hundreds of animals), we assessed *rme-2* using the short-reversal assay, which is scored on individual animals; we found that *rme-2* mutants are defective for P11-induced short reversals (Fig. 1I). Together, our data suggest that SID-1 does not function in any of the “establishment” steps in the intestine or germline upstream of Cer1 VLP capsid formation and packaging; instead, RME-2 rather than SID-1 appears to act at the germline entry step, and SID-1 acts at a point downstream of the intestine and germline.

### SID-1 does not function in neurons to mediate sRNA-induced learning

It has been previously shown that expression of *sid-1* is barely or undetectable in neurons^13,26,27^; our pan-neuronal^28^ and adult neuronal single-nucleus RNAseq data from adult neurons^29^ supports this conclusion, with expression levels less than 0.04 counts/cell (Fig. 2A; Fig. S1A). Consistent with those previous findings, reduction of *sid-1* function in neurons via a pan-neuronally expressed dsRNA construct (*Prgef-1::sid-1(RNAi*) did not show any defects in naive or learned avoidance (Fig. 2B), suggesting that SID-1 does not function in the neurons to mediate sRNA-induced learning. Together, our results suggest that SID-1’s true site of function in pathogen-induced learned avoidance is not the intestine, germline, or neurons.

**Figure 2:**
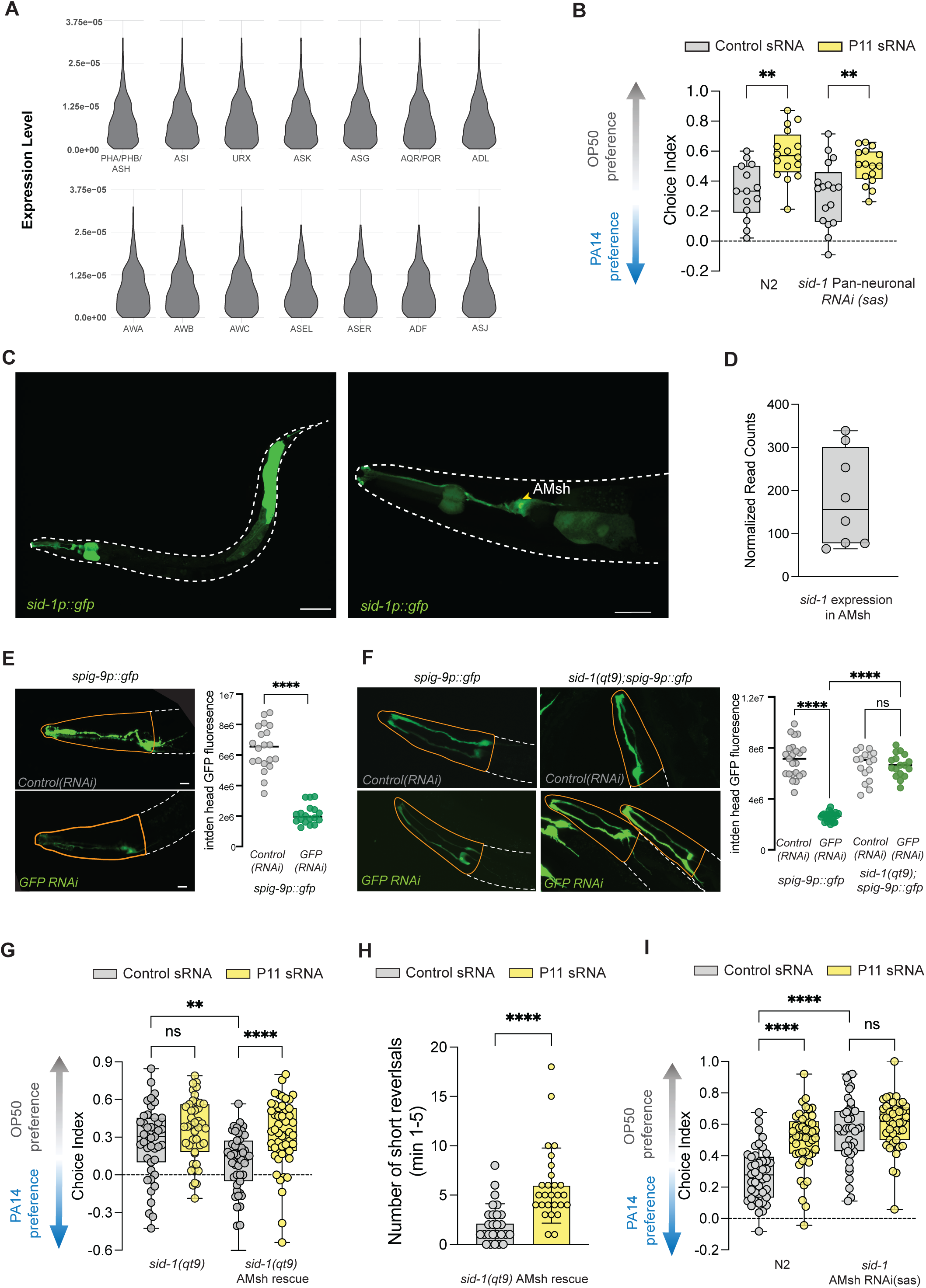
SID-1 is expressed and functions in the AMsh. A. Violin plots of *sid-1* expression in neurons from single-nucleus RNA sequencing data^29^; *sid-1* expression is low to undetectable in most neurons. B. Loss of *sid-1* in neurons (via pan-neuronal sid-1 RNAi) does not affect learned pathogen avoidance with P11 training. C. *sid-1p::gfp* is expressed in the pharynx and intestine, as previously reported. Scale bar represents 100µm (*left*). *sid-1p::gfp* is expressed in the amphid sheath glial cells. Scale bar represents 10µm (*right*). D. *sid-1* expression read counts in the amphid sheath glia cells isolated using FACs. Each dot represents expression data from an independent replicate. n = 7 E. Amphid sheath glial cells are sensitive to RNA interference: RNAi shows knockdown of GFP in amphid sheath glia. Representative fluorescent images of C. elegans heads in which *spig-9::gfp* is expressed. Scale bar represents 10µm. (left) Scatter plot shows quantification of the indent for the ROI (marked in red). Three independent biological replicates were quantified (right). \*\*\**\*p<*0.0001(Welsh’s t-test). F. Loss of *sid-1* from AMsh prevents RNAi of GFP. Representative fluorescent images of C. elegans heads in which *spig-9::gfp* in wild-type and *sid-1* animals is expressed. Scale bar represents 10µm (*left*). Scatter plot shows quantification of the indent for the ROI (marked in red). Three independent biological replicates were quantified (*right*). \*\*\*\**p<*0.0001(Two-way ANOVA with Tukey’s multiple comparison’s test). G. AMsh-specific sid-1 rescue in *sid-1(qt9)* mutant background restores naive behavior and learned pathogen avoidance with P11 training. H. AMsh-specific *sid-1* rescue in *sid-1(qt9)* mutant background restores P11-induced short reversals. I. AMsh-specific *sid-1* knockdown (via dsRNA of *sid-1* expressed in the AMsh) increases naive avoidance and prevents learned pathogen avoidance with P11 training. For choice assays each dot represents an assay plate of 20-156 worms, average = 64 worms /plate, average of 14 plates/genotype/training condition; n ≥ 3 biological replicates. Boxplots: center line represents median; box range denotes the 25th to 75th percentile; whiskers represent minimum to maximum. Two-way ANOVA with Tukey’s multiple comparison’s test (B, H-J). \*\*\*\**p<*0.0001, \*\*\**p<*0.001, \*\**p<*0.01, \**p<*0.05, ns = not significant. Scatter plot: each dot represents a neuron; center line denotes median.

### sid-1 is expressed in AMsh glia and enables RNA interference in the AMsh glia

To identify the tissue in which SID-1 functions in sRNA-induced pathogen avoidance, we first re-examined the expression pattern of *sid-1* using a promoter-GFP *sid-1p::gfp* strain. Although expression of *sid-1p::gfp* in previously identified tissues, including intestine and pharynx, was confirmed (Fig. 2C), we also noticed that *sid-1* is also prominently expressed in the head region; further examination revealed that *sid-1* is expressed specifically in the amphid sheath glial cell (AMsh) (Fig. 2D), a nervous system cell that envelops several amphid neurons. This was surprising, as expression of *sid-1* in the nervous system had not been previously reported. FACS sorting of the AMsh and subsequent RNA-seq confirm that *sid-1* is significantly expressed in the AMsh (Fig. 2E).

Moreover, we found that RNA interference, which is not functional in neurons due to the lack of neuronal *sid-1* expression^27^, is functional in the amphid sheath: expression of GFP in the AMsh glia in an otherwise wild-type background is susceptible to RNAi of GFP (Fig. 2F), unlike the well-documented resistance to RNAi in *C. elegans* neurons^27^, and loss of *sid-1* renders the animals defective for RNAi of GFP in the AMsh (Fig. 2G). Thus, AMsh glial cells, unlike neurons, are susceptible to RNA interference, and SID-1 is required for this ability.

### *sid-1* expression in amphid sheath glia is both necessary and sufficient for sRNA-induced learned pathogen avoidance

Next, we investigated the role of SID-1 in the AMsh in P11-induced learned avoidance. First, we found that expression of *sid-1* in the amphid sheath glial cell of a *sid-1* mutant rescues naive attraction to normal levels (Fig. 2H) and also rescued P11-induced learned avoidance (Fig. 2H), as well as P11-induced short reversals (Fig. 2I). Therefore, expression of *sid-1* specifically in the amphid sheath glia is sufficient to rescue both the naive and sRNA-induced phenotypes of *sid-1* mutants.

Perhaps more importantly, loss of *sid-1* specifically in the AMsh both increased naive avoidance and prevented P11-induced learned avoidance (Fig. 2J), phenocopying the behaviors of *sid-1* mutants, in contrast to the lack of effect of *sid-1* knockdown in neurons (Fig. 2B). Together, our results show that *sid-1* expression in amphid sheath glia, and only in amphid sheath glia, is both necessary and sufficient for P11-induced learned avoidance. Thus, the amphid sheath glial cells appear to be the true and sole site of SID-1 function in mediation of naive attraction to and environmental sRNA-induced learned avoidance of PA14.

### *SID-1* loss does not compromise general AMsh development, morphology, or sensory function

Because the AMsh ensheathes multiple amphid sensory neurons and supports several SID-1-independent behaviors, we asked whether loss of *sid-1* compromises the AMsh generally, which would confound interpretation of its role in P11-induced learning. We found that gross AMsh morphology (Fig. 2E) and function were not affected by *sid-1* loss of function: baseline, untrained *daf-7p::gfp* expression in the AMsh-ensheathed ASI neuron was unaffected by loss of *sid-1* (Fig. 1C, D), consistent with normal AMsh-ASI structural association in the absence of training. Finally, chemotaxis to pyrazine, an AWA-dependent volatile attractant^30,31^, was normal in *sid-1* mutants (Fig. S2A), indicating that sensory function is not grossly disturbed by loss of *sid-1* function. Together, these results indicate that SID-1 is dispensable for the AMsh’s structural integrity and for its general support of sensory behavior, and that its specific requirement in P11-induced learned avoidance reflects a discrete molecular function rather than a general defect in AMsh health or development.

### SID-1’s RNA binding domain is required for naive and learned avoidance

Next, we tested whether SID-1 is likely to interact with RNAs from the germline. We previously found that the germline-expressed Cer1 Ty3/Gypsy retrotransposon^32^ is required for sRNA-mediated learned pathogen avoidance through its formation of virus-like particles that contain RNA^8^, and that Cer1 acts not in the germline, but rather in the communication between the germline and neurons^8^. Therefore, SID-1 in the amphid sheath glia may interact with Cer1 delivered-RNA. To test this hypothesis, we mutated the RNA binding residues of the RNA binding domain of SID-1^18,23^(Fig. 3A); we expressed this form of SID-1 specifically in the amphid sheath glia of the otherwise *sid-1(qt9)* mutant background strain. We found that this RNA-binding mutant (RBD) form of SID-1 was unable to rescue naive or learned avoidance (Fig. 3B, in contrast with the rescue exhibited by intact SID-1 (Fig. 2F), suggesting that the RNA binding domain of SID-1 is critical.

**Figure 3:**
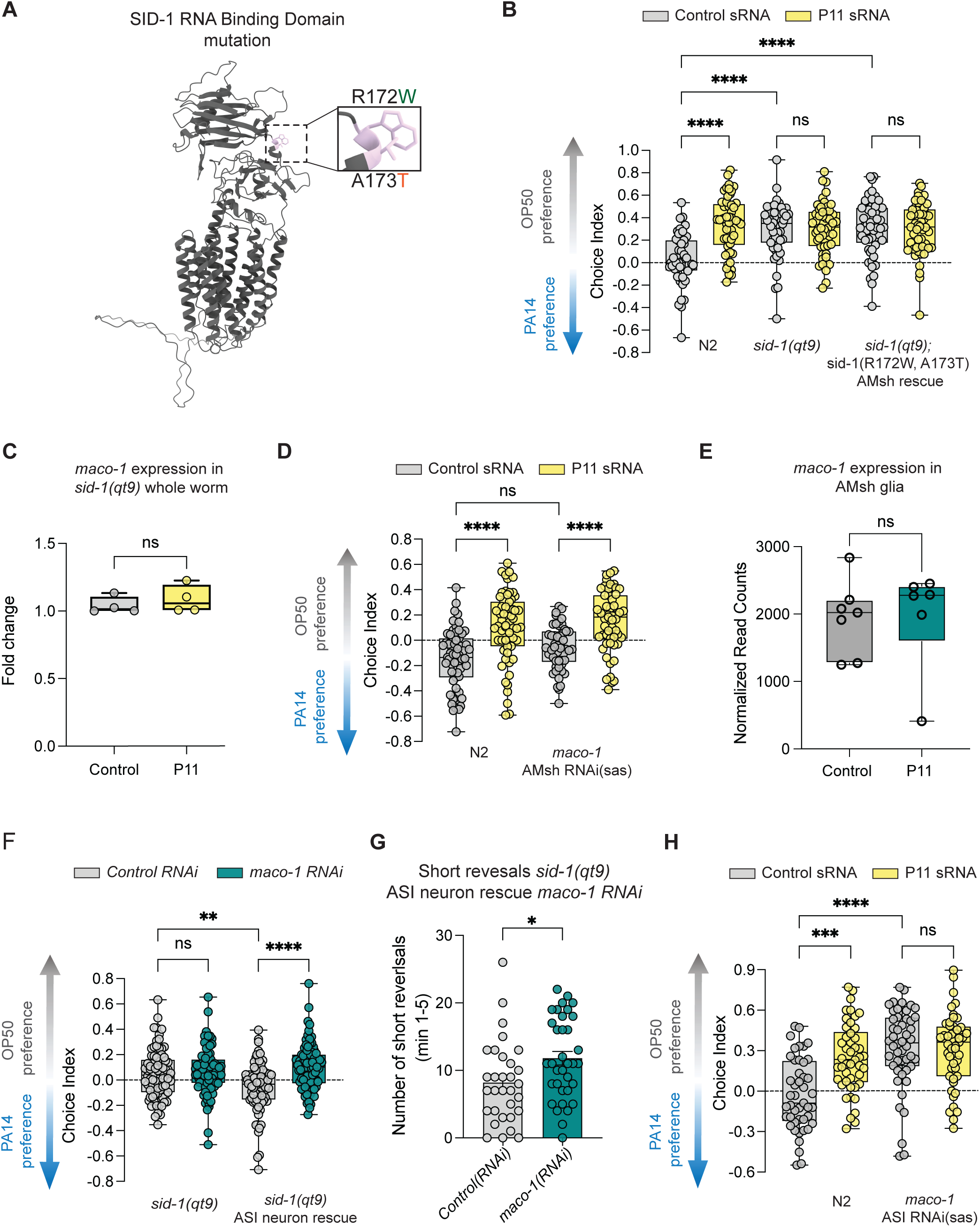
*maco-1* is not altered in the AMsh, but rather in the ASI neuron. A. Mutations in R172 and A173 of SID-1 to Tryptophan and Threonine respectively, alter RNA binding^18^. B. The RNA binding domain of SID-1 expressed in the AMsh is required for rescue of naïve choice and learned behavior with P11 sRNA training. C. *sid-1* is required for changes in *maco-1* levels upon P11 training. Boxplots showing the relative mRNA levels of *maco-1* ± P11 training by qPCR. Each dot represents a biological replicate, *n = 4*. ns (not significant) *p =* 0.5726 t-test. D. Reduction of *maco-1* specifically in the AMsh did not induce high naive avoidance of PA14, and the animals still earned avoidance upon training with P11. E. RNA-sequencing of FACS-sorted AMsh cells ± P11 training revealed that *maco-1* levels are not altered in amphid sheath glia. Each dot represents a biological replicate, n = 7. *p =* 0.9444 Welsh’s t-test. amphid sheath glia. Each dot represents a biological replicate, n = 7. *p =* 0.9444 Welsh’s t-test. F. *sid-1(qt9)* mutants with *sid-1* rescued only in the ASI exposed to adult-only *maco-1* RNAi (L4 to day 1 adult) *maco-1* RNAi bacteria have high naive avoidance of PA14 G. Bar plot showing *sid-1(qt9)* mutants with *sid-1* rescued only in the ASI treated exposed to adult-only *maco-1* RNAi (L4 to day 1 adult) bacteria exhibit increased short reversals. *n = 3* independent replicates. *p<0.05 Welsh’s t-test. H. Reduction of *maco-1* specifically in the ASI (via dsRNA of maco-1 expressed only in the ASI) increases naive avoidance and prevents P11-induced learned avoidance. For choice assays each dot represents an assay plate of 26-220 worms, average = 97 worms /plate, average of 20 plates/genotype/training condition; *n* ≥ 3 biological replicates. Boxplots: center line represents median; box range denotes the 25th to 75th percentile; whiskers represent minimum to maximum. Two-way ANOVA with Tukey’s multiple comparison’s test (B, D, F, H). \*\*\*\**p<*0.0001, \*\*\**p<*0.001, \*\**p<*0.01, \**p<*0.05, ns = not significant. Scatter plot: each dot represents a neuron center line denotes median.

### maco-1 is not targeted directly in the amphid sheath glia

The P11 sRNA has a 17-nucleotide perfect match to the macoilin homolog *maco-1*^6^, which is downregulated in P11 sRNA trained wild-type worms^4–6^. We found that *maco-1* levels do not further decrease upon P11 training in *sid-1(qt9)* animals (Fig. 3C).

We reasoned that if SID-1 delivers RNA containing the 17nt match to the target cell, *maco-1* should be downregulated in the cell that receives the P11 sRNA fragment products. Therefore, we tested whether knockdown of *maco-1* specifically in the AMsh phenocopies P11 training or *sid-1* knockdown in the amphid sheath glia; however, reduction of *maco-1* specifically in the AMsh did not induce high naive avoidance of PA14, and the animals still learned avoidance upon training with P11 (Fig. 3D); thus, *maco-1* does not appear to be targeted in the amphid sheath glia, counter to our expectations. To confirm these results, we FACS isolated AMsh cells ± P11 training; *maco-1* levels were not altered upon P11 training (Fig. 3E), consistent with our behavioral results. Together, these data suggest that while SID-1 is required in the amphid sheath glia, AMsh is not the site of *maco-1* targeting by P11 RNA products carried from the germline by Cer1.

### maco-1 in the ASI neuron is required for pathogen avoidance upon P11 training

We previously found that the ASI neuron, where *daf-7p::gfp* increases upon P11 training, is required for learned avoidance upon PA14 training^6^. Therefore, we next asked whether *maco-1* is targeted by P11-related RNA in the ASI upon P11 training. Although *sid-1* is not required in neurons for sRNA-induced pathogen learning (Fig. 2B), we found that artificial expression of *sid-1* in neurons, which is used standardly to allow neuron-specific RNAi^27^, is sufficient to suppress the high naive avoidance and allow rescue of the P11-induced learned avoidance (Fig. S3A). Similarly, expression of *sid-1* only in the ASI neuron allowed P11-induced learning, and rescued P11-induced short reversals (Fig. S3B-C). We therefore used the same principle to enable RNAi specifically in the ASI neuron to test whether it is the site of *maco-1* knockdown. Indeed, *maco-1* RNAi knockdown only in the ASI caused high naive avoidance of PA14 (Fig. 3F) and increased short reversals (Fig. 3G), similar to *maco-1* mutants.

To complement this approach, we expressed *maco-1* dsRNA in the ASI in otherwise wild-type animals and found that naive avoidance is increased, and these animals cannot learn on P11 (Fig. 3H). These effects on ASI by *maco-1* decreases contrast with the lack of effect by *maco-1* RNAi treatment in the amphid sheath (Fig. 3B).

Together, our data suggest that the cellular target of *maco-1* reduction by the P11 sRNA products is the ASI neuron. That is, normally, SID-1 function in the AMsh is required for *maco-1* reduction in the ASI upon P11 training, suggesting a non-autonomous role for SID-1 in its effect on maco-1 expression and learned avoidance behavior.

### Endocytic and secretory pathway components are required in the AMsh for P11-induced learned avoidance

Because Cer1 delivers RNA to the AMsh, but its target is *maco-1* in the ASI rather than the AMsh itself, we asked whether intracellular trafficking machinery in the AMsh is required for these ASI-driven phenotypes. Clathrin-mediated endocytosis is a key mechanism for internalizing material from the cell surface; clathrin assembles into coated pits on the inner face of the plasma membrane^33,34^. Adaptor proteins such as AP180 help recruit clathrin and drive membrane curvature, shaping the pit as it forms^35,36^. Dynamin then mediates scission, pinching off the budding pit to release a clathrin-coated vesicle into the cell^33,34,36^. A separate, clathrin-independent route uses caveolae, which mediates endocytosis in parallel to the clathrin-dependent pathway^37^. These components have also been implicated in transcytosis, the process of shuttling endocytosed cargo through one cell to another^34,38^. Therefore, we wondered if these endo/transcytosis components might play a role in the transfer of information through the amphid sheath glia to the ASI.

Using our amphid sheath glia-specific RNAi strain, we knocked down the clathrin heavy chain (*chc-1*) briefly from late L4-adulthood by RNAi (to avoid disrupting amphid sheath glia or neuronal development) and carried out P11 training. We found that briefly reducing clathrin specifically in the AMsh glia both increased naive avoidance and abolished P11-induced avoidance (Fig. 4A), reminiscent of *sid-1* and *maco-1* mutants. Brief reduction of clathrin adaptor protein AP180/*unc-11*, caveolin (*cav-1*), and dynamin (*dyn-1*) specifically in amphid sheath glial cells also abolished P11-induced learning (Fig. 4B-D). (We cannot interpret the lack of an effect on naive avoidance in these experiments, as RNAi may have been too brief or incomplete.) These data indicate that endocytic and/or secretory trafficking in the AMsh is necessary for P11-mediated learned avoidance.

**Figure 4:**
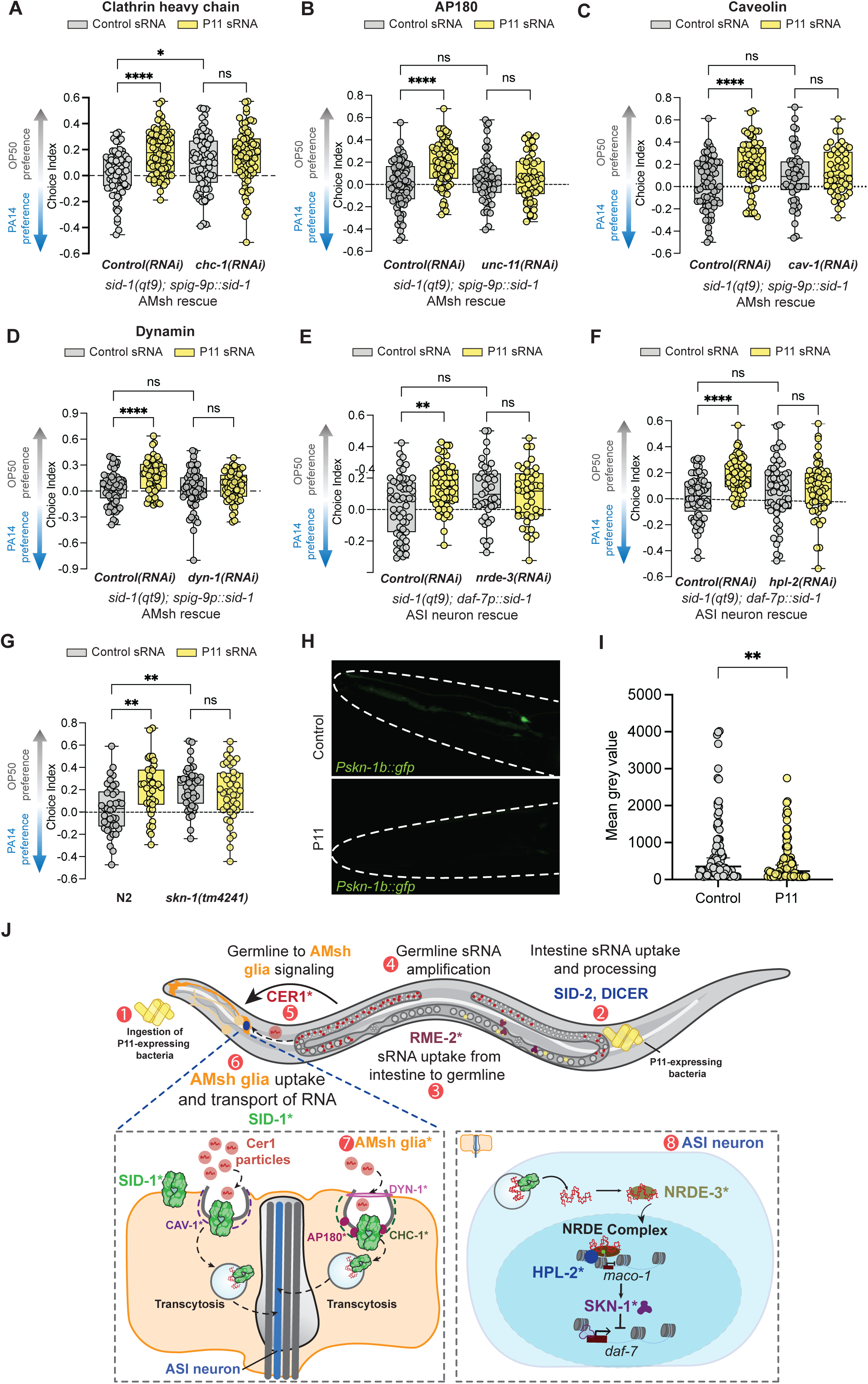
AMsh-to-ASI RNA communication is required for P11-induced learned avoidance. A. Brief reduction of clathrin heavy chain (*chc-1*) with RNAi in the amphid sheath glia increases naïve avoidance and prevents P11-induced learned avoidance. B. Brief reduction of unc-11/AP180 (*unc-11*) with RNAi in the amphid sheath glia prevents P11-induced learned avoidance. C. Brief reduction caveolin (*cav-1*) with RNAi in the amphid sheath glia prevents P11-induced learned avoidance. D. Brief reduction of dynamin (*dyn-1*) with RNAi in the amphid sheath glia prevents P11-induced learned avoidance. E. ASI-specific RNAi of *nrde-3* reduces P11-mediated learned pathogen avoidance. F. ASI-specific RNAi of *hpl-2* reduces P11-mediated learned pathogen avoidance. G. *skn-1(tm4241)* mutants exhibit high naïve avoidance of PA14 and cannot learn avoidance with P11. H. *Pskn-1b::gfp* is expressed in the ASI neuron, as previously reported^45,46,50^, but expression is reduced upon treatment with P11 E. coli. I. Quantification of expression levels on control and P11 E. coli. J. Model of SID-1’s role in the amphid sheath glia to carry RNA signals from the germline to the ASI. *Denotes findings first established in this study. For choice assays each dot represents an assay plate of 20-152 worms, average = 51 worms /plate, average of 22 plates/genotype/training condition; n ≥ 3 biological replicates. Boxplots: center line represents median; box range denotes the 25th to 75th percentile; whiskers represent minimum to maximum. Two-way ANOVA with Tukey’s multiple comparison’s test (A-G). \*\*\*\**p<*0.0001, \*\*\**p<*0.001, \*\**p<*0.01, \**p<*0.05, ns = not significant. Scatter plot: each dot represents a neuron center line denotes median.

### RNA-interacting proteins are required in the ASI

If RNA is transferred from the AMsh to the ASI, we hypothesize that the RNA must have interactors in the target cell, the ASI neuron, in order for *maco-1* levels to be reduced. While many RNA-binding chromatin modifiers are expected to function in the germline, previous studies showed that *nrde-3* and *hpl-2* can act in specific neurons to regulate adaptation through endo-siRNAs^39^. NRDE-3 (nuclear RNAi defective-3) carries RNAs from the cytoplasm to the nucleus^40,41^, and HPL-2 promotes RNA-directed transcriptional gene silencing^42,43^. We had previously found that *nrde-3*^7^ and *hpl-2* ^3,6^ are required for PA14- and P11-induced learned avoidance, but because those experiments were done in whole animal mutants, the site of function was not determined. To test the cell-specific requirement for these genes, we again used the mutant expressing *sid-1* only in the ASI to enable ASI-specific knockdown of these genes by treating the worms with bacteria expressing double-stranded RNA of *nrde-3* and *hpl-2.* We found that reduction of *nrde-3* and *hpl-2* specifically in the ASI each inhibit P11-mediated learning (Fig. 4D, E). Interestingly, the loss of these genes does not alter naive avoidance of PA14, which is consistent with the loss of the ability to reduce *maco-1* levels, which would render animals unable to learn from P11 training but unchanged in their naive avoidance state. Our data are the first to suggest that these same components that function in endo-siRNA regulation are critical for the interpretation of signals that originate from exogenous environmental RNA signals.

### SKN-1 is required for naive and learned avoidance

The Nrf2-related SKN-1 transcription factor plays a role in xenobiotic resistance, stress response, longevity, and pathogenic responses^44–46^, and is known to be expressed in the ASI^45,47^, which we confirmed (Fig. S3D). SKN-1 inhibits the expression of *daf-7*, which we previously found is correlated with PA14 and P11 training^3,6^ (Fig. 1D). Therefore, we tested whether SKN-1 plays a role in the regulation of naive or P11-mediated learned avoidance. *skn-1(tm4241)* mutants, like *maco-1* and *sid-1* mutants, displayed high naive avoidance and could not learn to further avoid PA14 (Fig. 4G), suggesting that SKN-1 function is key for this pathway. We then asked whether skn-1 expression is regulated by the PA14 learned avoidance pathway; indeed, *Pskn-1b::gfp* expression is decreased in animals trained on P11 E. coli (Fig. 4H, I).

### Model of P11-induced learned avoidance

The simplest model that accounts for all our data (Fig. 4J) involves *C. elegans* ingesting *Pseudomonas aeruginosa* PA14’s small RNA, P11, into the worm’s intestine via SID-2, processing by Dicer in the intestine, and transport into the germline by RME-2. There it is processed further by RNA-dependent polymerases^3,6^ then carried by Cer1 VLPs to the amphid sheath glial cells. At the AMsh glia, the RNA carried by Cer1 meets SID-1, and endo/transcytosis components then are required for transport to the ASI neurons, where NRDE-3 interacts with the RNA, resulting in HPL-2-dependent silencing of *maco-1* transcription. Finally, reduction of SKN-1, which is downregulated upon P11 training and subsequent maco-1 downregulation, allows an increase in daf-7 expression, resulting in the switch from naïve attraction to avoidance of PA14.

## DISCUSSION

Learning to avoid danger is key for survival. *C. elegans* learns to avoid pathogenic *Pseudomonas* through interpretation of bacterial small RNAs, passing this learned information on for four generations and horizontally to kin. While the RNA interference components Dicer and SID-2 are required in the intestine, the tissue(s) in which the dsRNA transporter SID-1 functions was unclear. Our experiments revealed that SID-1 does not act in the germline or intestine; instead, SID-1 function in amphid sheath glia (AMsh) is both necessary and sufficient for learned avoidance of *Pseudomonas aeruginosa* (PA14) via its sRNA, P11. P11’s target, *maco-1*, is not downregulated in the AMsh, but rather in the ASI neuron, which is enveloped by the AMsh glial cell - a process requiring endo/transcytosis components. Our data suggest that RNA is transported from the germline to the AMsh glia, where SID-1 enables uptake and transfer to the ASI neuron and subsequent *maco-1* downregulation, resulting in avoidance of PA14. These data suggest a new role for SID-1 in endocytic RNA uptake into the amphid sheath glia, and identify the AMsh as an RNA filter gating germline-to-neuron communication of environmental small RNA signals, an invertebrate parallel to the mammalian blood-brain barrier.

Our *in vivo* results resolve several important components of the SID-1 model that recent structural work could not address. We uncovered a new role for SID-1 in an unexpected cell of the nervous system: SID-1 does not act in the intestine or germline to regulate environmental RNA-dependent behavior, but instead functions in a single, non-neuronal cell type in the nervous system, the amphid sheath glia. Together with the emerging structural picture of SID-1/SIDT1/SIDT2 as endocytic dsRNA receptors, rather than passive channels, that are capable of binding small RNA substrates substantially shorter than previously proposed, our data complete a broader revision of two-decade-old assumptions about where and how this RNA transporter functions. We propose that the amphid sheath glia constitute an RNA filter that gates which environmentally derived, germline-amplified RNA signals reach the nervous system through Cer1 viral-like capsids. Neurons in *C. elegans* are well known to be refractory to RNA interference^27^, a property generally interpreted as protective, due to their lack of SID-1 expression. Our data show that this protection is likely not an absolute exclusion but instead a gated passage through a single, RNAi-competent intermediary: the AMsh, unlike neurons, appreciably expresses SID-1and is fully RNAi-sensitive. The AMsh therefore constitutes the sole licensed entry point through which an otherwise-excluded class of RNA signals is permitted to influence neuronal gene expression and behavior, with SID-1 restricting RNA uptake to this cell rather than allowing RNA to be delivered to neurons directly. This is, to our knowledge, the first description of a dedicated glial cell type acting as an obligate RNA gateway between an animal’s external chemosensory environment and its nervous system.

Our data indicate that the AMsh is a required but transient location for this RNA signal rather than its site of action: *maco-1* is unaltered in the AMsh itself, and P11-dependent silencing occurs specifically in the ASI. Our data support a model in which SID-1 mediates endocytic uptake of Cer1-delivered RNA at the AMsh followed by transfer to the adjacent ASI neuron, consistent with the shared requirements for clathrin, AP180/UNC-11, dynamin, and caveolin, which have been previously implicated in transcytosis^34,38^. SID-1 function in the AMsh is required to relay the RNA signal, and the ASI is the site where that RNA signal is ultimately interpreted through interactions with NRDE-3 and HPL-2, culminating in regulation of *maco-1* expression and subsequent activity of the SKN-1 transcription factor to regulate the TGF-beta ligand DAF-7. The requirement for endocytic/secretory machinery in the AMsh, the RNA-binding-dependent requirement for SID-1’s function there, the strict spatial restriction of rescue and loss-of-function effects to the AMsh (*sid-1*) and ASI (*maco-1*), and the demonstration that these effects are not attributable to a general loss of AMsh integrity or sensory function all suggest a role for the amphid sheath glial cells in transcytosis of an RNA signal to neurons.

Our data also at least partially resolve a previously confusing observation: the phenotypes of *sid-1* mutants (i.e., high naïve avoidance and inability to learn P11-induced avoidance) more closely resemble *maco-1* mutants than those of intestinal gene mutants (*sid-2*, *dicer*) or germline gene mutants, as *sid-1*, Cer1, and *maco-1* mutants already exhibit high naive avoidance of PA14 prior to any training^6,8^, while the others only lack P11-induced learned avoidance. Here we found that rescue of *sid-1* expression solely in the AMsh restores the low naive avoidance and ability to learn upon P11 training; furthermore, loss of *sid-1* specifically in the AMsh induces high naive avoidance, reduction of at least some endo/transcytosis genes in the AMsh cause high naive avoidance, and *maco-1* reduction specifically in the ASI induces high naive avoidance. Similarly, loss of function of *skn-1*, a transcription factor that regulates ASI function, also results in high naive avoidance behavior. Thus, mutants of every component that acts downstream of the germline in packaging and transporting the small RNA to the nervous system to regulate *maco-1* levels all share this high naive avoidance phenotype, which appears to be a sign of constitutively low *maco-1* levels. These data support a model in which the loss of *sid-1* in the AMsh results in low *maco-1* in the ASI, resulting in an avoidance of *P. aeruginosa* without prior exposure, as *maco-1* acts as a rheostat for avoidance activity. However, these results also suggest that under normal conditions on non-pathogenic bacteria, an unidentified, perhaps endogenous signal that may keep *maco-1* levels high must also be present, as mutants in the early part of the pathway (intestine and germline) do not display altered naive avoidance.

Our data also address a more general question: how can circulating RNAs, even those in extracellular vesicles, be delivered to their target cells? The RNA gating role of SID-1 in the AMsh evokes an analogy to the mammalian blood-brain barrier, where astrocytes serve as selective intermediaries between systemic circulation and neurons, taking up circulating metabolites, ions, and macromolecules rather than allowing direct neuronal access, a relationship that has been described as a protective physiological filter for the nervous system. The AMsh occupies a directly analogous position with respect to RNA, interposed between an external signal source and the neurons that respond to the RNA. The parallel extends to the underlying molecular machinery: SID-1 and its mammalian orthologs SIDT1/SIDT2 share the same core biochemistry of endolysosomal dsRNA uptake and pH-dependent small RNA binding. We propose that the AMsh represents an invertebrate, single-cell-type instance of the same general design principle that the blood-brain barrier exemplifies in mammals: a glial barrier that does not simply exclude environmental nucleic acid signals from the nervous system but actively receives and relays a subset of them onward, thereby determining which external information the nervous system is exposed to at all. The relative simplicity and genetic tractability of the *C. elegans* nervous system make the AMsh a uniquely accessible system in which to dissect a process that in mammals is distributed across a far more complex and less experimentally tractable barrier.

## Supporting information

Table S1 Strains

Table S2 raw data

## RESOURCE AVAILABILITY

All data needed to evaluate the conclusions presented in this paper are available in the main text or the supplementary materials. BioProject # PRJNA1521498. Further information and requests for resources and reagents should be directed to.

## ACKNOWLEDGMENTS

We thank the *Caenorhabditis* Genetics Center for *C. elegans* strains. We also thank W. Wang, J. Arly Volmar, and J. Miller (Genomics Core Facility, Princeton University) and C. DeCoste and the Molecular Biology Flow Cytometry Resource Facility, which is partially supported by the Rutgers Cancer Institute of New Jersey NCI-CCSG P30CA072720-5921, and J. Ferencz for assistance; and the Murphy lab for discussion and feedback. C.T.M. is the Director of the Lewis Sigler Institute for Integrative Genomics and the Simons Foundation for Plasticity in the Aging Brain. This work was supported by a Pioneer Award to C.T.M. (NIGMS DP1GM119167), a Simons Foundation grant to C.T.M. (AN-NC-AB-Research-00811235SPI), the Princeton Presidential Postdoctoral Fellowship to A.B., Ford Foundation Fellowship R.S., and T32GM148739 for support of K.N.

## AUTHOR CONTRIBUTIONS

Conceptualization: A.K.B., R.K., and C.T.M.; methodology: A.K.B., X.L., R.K., R.S., K.N., L.S., and C.T.M.; Investigation: A.K.B., X.L., R.K., R.S., K.N., and L.S.; writing – original draft: A.K.B. and C.T.M.; writing – review & editing: A.K.B., R.K., and C.T.M.; funding acquisition: C.T.M.; supervision: C.T.M.

## DECLARATION OF INTERESTS

The authors declare no competing interests.

## SUPPLEMENTAL INFORMATION

Document S1. Supplemental Figures S1-S3

Data S1. Method on strain generation

Data S2. Source data on all figures

## STAR METHODS

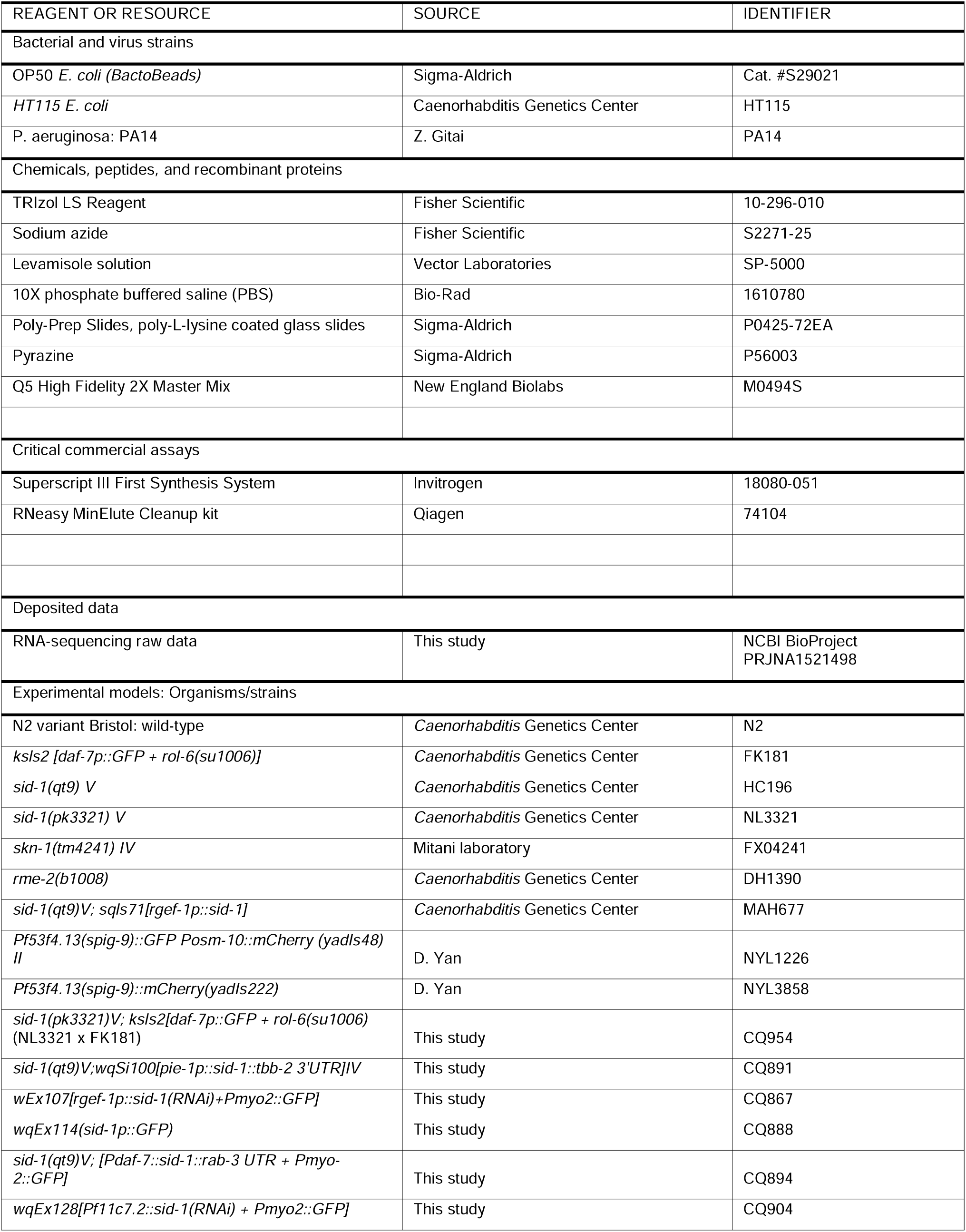
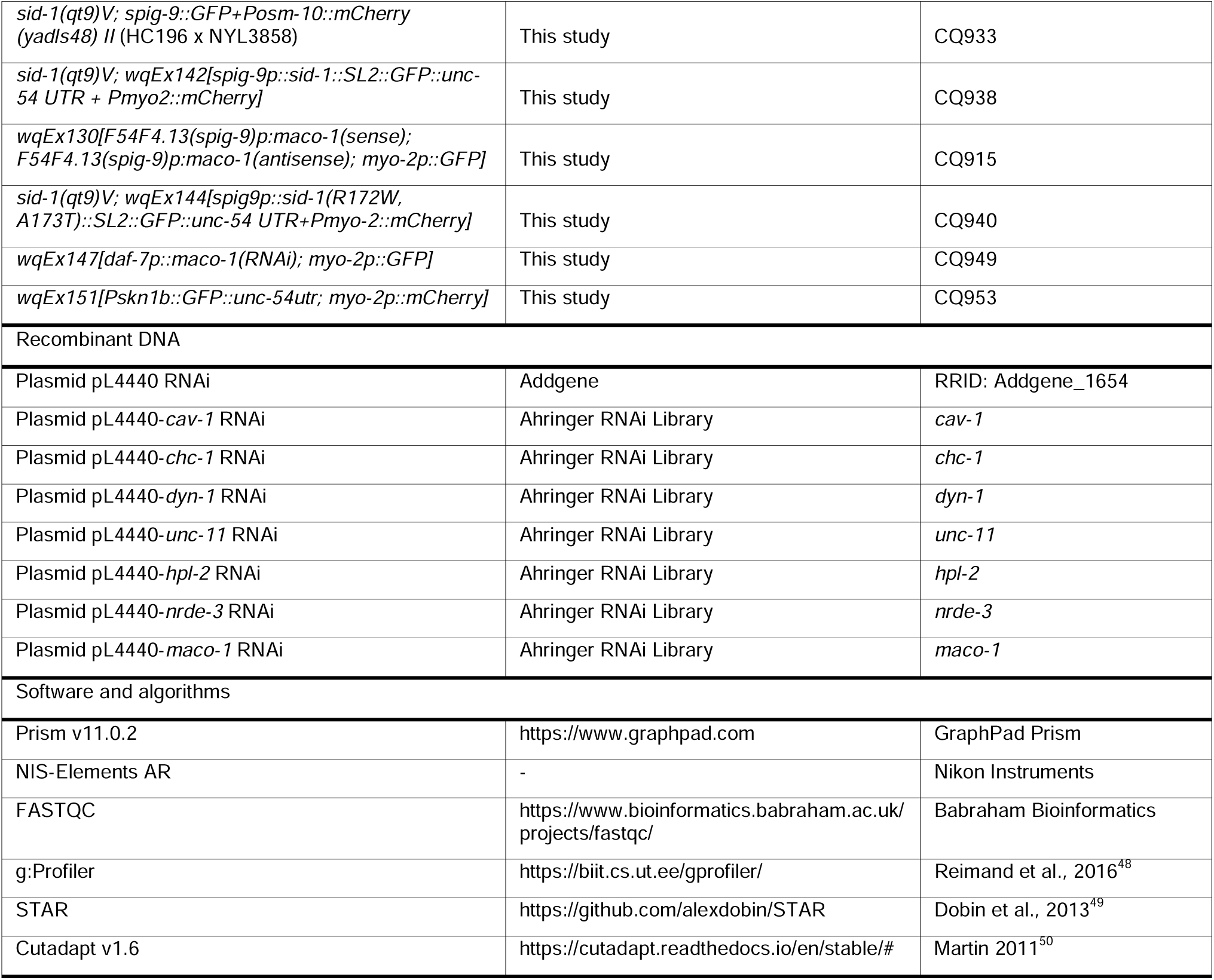

### EXPERIMENTAL MODEL AND SUBJECT DETAILS

#### C. elegans strains

The list of all strains used, along with methods for generation, for this study can be found in Data S1.

### METHOD DETAILS

#### General maintenance of *C. elegans* strains

Worm strains were maintained at 20°C on high growth medium (HGM) plates (3 g/L NaCl, 20g/L bactopeptone, 30 g/L bacto-agar in distilled water, with 4mL/L cholesterol [5 mg/mL in ethanol], 1 mL/L, 1 M CaCl_2_, 1 ml/L MgSO_4_, and 25 mL/L 1 M potassium phosphate [pH 6.0] added to molten agar after autoclaving) seeded with *E. coli* OP50, using standard methods. Plates were supplemented with carbenicillin and tetracycline for RNAi bacteria growth, and kanamycin for P11 expressing *E. coli*.

#### Bacterial strains

*E. coli* OP50 was obtained from the *C. elegans* Genetics Center (CGC) and *P. aeruginosa* PA14 was a gift from Z. Gitai’s laboratory. *E. coli* expressing the P11 sRNA was cultured overnight in Luria broth supplemented with 50µg/mL kanamycin.

RNAi strains: All RNAi strains are from the *C. elegans* RNAi Collection (Ahringer library). These include *L4440* RNAi control, *maco-1* RNAi, *chc-1* RNAi, *unc-11* RNAi, *dyn-1* RNAi, *cav-1* RNAi, *nrde-3* RNAi and *hpl-2* RNAi strains.

#### Cultivation of bacterial strains

*E. coli* OP50, *P. aeruginosa* PA14, and *E. coli* expressing the P11 sRNA were cultured overnight (14–16 hours) in LB (10 g/L tryptone, 5 g/L yeast extract, 10 g/L NaCl) with shaking (250 rpm) at 37°C. *E. coli* strains carrying the P11 expression plasmid were cultured in LB supplemented with kanamycin (50 µg/mL). RNAi bacteria (tissue-specific and whole-animal) were grown in LB supplemented with carbenicillin (100 µg/mL) and tetracycline (12.5 µg/mL) at 37°C, overnight for (approximately 16 hours) with shaking.

#### sRNA Training

Training plates were prepared by seeding HG/kanamycin plates with 800 µL of bacteria P11-expressing *E. coli* and incubating at 25°C for 48 hours to allow lawn growth. On the day of training plates were moved to room temperature to equilibrate for an hour before animals began training.

Young adult hermaphrodites were bleached to obtain eggs and placed on OP50 seeded HG plates. These worms were raised for 2 days at 20°C. Synchronized L4 animals were washed from OP50 plates with M9 and allowed to pellet. Approximately 35µL of pelleted animals were added to each training plate and incubated at 20°C for 24 hours.

#### Bacterial choice assay plate preparation

Overnight bacterial cultures were diluted in LB to an OD_600_ of 0.5, and 25 µL of each bacterial suspension was spotted onto opposite sides of a 6-cm NGM plate. Plates were incubated for 2 days at 25°C and equilibrated to room temperature (20°C) for 1 hour before the choice assay.

#### Aversive learning (choice) assay

On the day of the assay, choice plates were equilibrated to room temperature (20°C) for 1 hour. 1 µL of 1 M sodium azide was spotted onto each bacterial spot as a paralyzing agent. Worms were washed off training plates in M9, allowed to pellet by gravity, and washed 2–3 additional times to remove residual bacteria. Using a wide-orifice pipet tip, 5µL worms were spotted midway between the two bacterial lawns and allowed to choose for 1 hour at room temperature before counting. Plates with fewer than 20 worms (low N) or greater than 200 worms (to avoid clumping artifacts) were censored; an average of 67 worms/plate and average of 16 plates/genotype/training condition were used in each assay. Choice index = (# worms on OP50 − # worms on PA14)/(total # worms). Assays with animals with extrachromosomal arrays involved counting transgenic animals (using the Leica fluorescent microscope) and total animals on each plate.

For the F1 choice assay, 5µL of worms were spotted midway between the two bacterial lawns on either side to ensure there >20 transgenic animals on each choice plate, with an average of 45.6 per/plate, while avoiding clumping artifacts.

#### Short-reversal assay

Day 1 adult worms were picked individually onto unseeded 60-mm NGM plates equilibrated to room temperature. After 1 minute of equilibration, each worm was observed for 4 minutes, and the number of short reversals (<3 head swings) and long reversals (≥3 head swings) was recorded^5,51^.

#### Conditioned media / horizontal transfer assay

Progeny of P11-trained or control-trained mothers were washed off plates, allowed to settle, and washed again in M9 before removing supernatant. For every 50 µL of worm pellet, 300 µL of S-basal media was added. Worms were incubated in this suspension for 24 hours at 20°C to produce conditioned media. After 24 hours, plates were inspected to exclude any wells in which worms had died, exploded, or bagged. Conditioned media was then pooled and centrifuged for 10 minutes at 6500 x g. This was then passed through a 0.22-µm filter to clear bacteria. 200µl of conditioned media was added to an OP50 spot, dried, and used to train naive L4 worms for 24 hours before testing in the choice assay.

#### RNAi treatment

Experiments involving RNAi: RNAi bacteria were grown plates HG plates prepared with IPTG (1 mM) and carbenicillin, grown at room temperature for 24 hours. Plates then spotted with 0.1M IPTG and allowed to dry at room temperature for 1 hour before use.

RNAi experiments were performed using standard feeding RNAi method. Bacterial clones expressing the control construct (empty vector, pL4440) and the dsRNA we obtained from the Ahringer RNAi library and sequences were verified prior to use.

RNAi tissue-sensitive strains previously published or (MAH677 for pan-neuronal) or generated for this study uses *sid-1(qt9)* animals and rescues *sid-1* in specific tissues using tissue specific promoters (CQ894 *daf-7p* for ASI, CQ938 *spig-9p* for AMsh). 200µl of 0.1M IPTG was spotted unto plates with dsRNA-expressing bacteria from the Ahringer library grown on IPTG/carbenicillin plates and allowed to dry at room temperature for an hour. Synchronized worms were then placed on plates at L4 stage through adulthood to day 1. This is done by washing L4 animals off plates with M9 buffer and allowing them to settle to a pellet. After removing excess M9, 12-15µl of worms were placed unto RNAi plates for 24 hours before testing their behavior.

In experiments where the F1 generation was subject to RNAi treatment, P0 animals were trained with P11-expressing E. coli for 24 hours. Worms were then bleached unto OP50 plates and allowed to grow to L4 stage. F1 worms from P11 trained parents were washed off plates with OP50 and allowed to pellet. They were then washed two more times with M9. 12-15µl of worms L4 worms were placed onto RNAi plates for 24 hours prior to testing their behavior.

#### Imaging and image analysis

Worms were mounted on 2% agar pads and immobilized with 1 mM levamisole. Z-stack multichannel (DIC and GFP) images were acquired at 60× magnification on a confocal or widefield fluorescence microscope. Maximum intensity projections of head neurons were generated in FIJI for representative images.

For *daf-7p::gfp* worm quantification, mean fluorescence intensity was quantified by drawing a Bezier outline around each ASI cell body in the single z-plane containing the neuron body. This was done using NIS Elements software. Exposure times were matched between control and treatment groups imaged on the same day.

For *spig-9p::gfp* worm quantification we drew our region of interest (ROI) around the head of the worm just behind the pharyngeal bulb in the brightfield image. Using FIJI, we then measured the integrated intensity (intden) of the fluorescence in that region. Exposure times were matched between control and treatment groups imaged on the same day.

#### AMsh isolation via FACS sorting

Animals with mCherry expressed in the AMsh glia (*PF53F4.13(spig-9)::mCherry(yadIs222)*) were used to selectively sort AMsh glia following P11 training. Following 24 hours of P11 training worms, were prepared for cell isolation using previously described methods^28,52^ with some modifications. Briefly, day 1 synchronized worms were washed 5 times with M9 to remove excess bacteria. Worms were then treated with 750µl of lysis buffer (200nM DTT,0.25% SDS, 20nM HEPES pH 8.0, 3% sucrose). Worms incubated for 6.5 minutes at room temperature to break the worm cuticle. The worms were then washed 5 times with M9 and resuspended with 500µl of 20 mg/mL pronase from *Streptomyces griseus* (Sigma-Aldrich). Worms were incubated at room temperature (<20 minutes) with mechanical disruption every 2 minutes by pipetting up and down. When most of the worm bodies were dissociated, leaving only eggs and small debris, the lysis was stopped by adding ice-cold PBS buffer with 2% fetal bovine serum (Gibco). Prior to sorting, cell suspension was passed through a 5µm filter. Using a BD Biosciences FACSAria Fusion sorter mCherry positive cells were sorted directly into Trizol LS. Sorting gates were determined by comparing age-matched non-fluorescent cell suspension samples.

#### RNA extraction, library preparation, and sequencing

Using the Trizol-chloroform-isopropanol method we extracted RNA and then performed RNA cleanup using the RNeasy MinElute Cleanup Kit (Qiagen). RNA quality and quantification was assessed using the Agilent Bioanalyzer RNA Pico chip. Library generation was performed using 2 ng of RNA and the Ovation Revelo RNA-Seq library preparation kit with SPIABoost Probe Mix-*C. elegans* (Tecan Genomics). Library quality and concentration were measured using an Agilent Bioanalyzer DNA 12000 chip. Samples were then multiplexed and sequenced (paired-end) on NovaSeq S1 100nt Flowcell v1.5 (Illumina).

#### RNA-seq analysis

Transcripts were processed as previously described^28,52^. Sequencing read quality was evaluated using FASTQC. Illumina universal adaptors were removed using Cutadapt v1.6. Trimmed reads were then aligned to the *C. elegans* reference genome, ce11, using STAR. Aligned reads were assembled htseq-counts and the expression levels of all transcripts were estimated. Differential gene expression was determined using DESeq2. Genes with an adjusted *p-value* 0.05 considered significantly differentially expressed. Overrepresentation of GO pathways among significantly differentially expressed genes was performed with g:Profiler (https://biit.cs.ut.ee/gprofiler/gost). Data submitted as BioProject number PRJNA1521498.

#### Whole worm RNA isolation

Total RNA isolated for *C. elegans* following 24-hour P11 exposure (described in sRNA Training section). Adult (P0) worms were washed from plates with M9 and allowed to pellet. Excess bacteria was removed by washing the worms 2X with M9. Worm pellets were crushed in liquid nitrogen and transferred to an appropriate volume of Trizol LS (100µl of worm pellet in 900µl of Trizol LS). Total RNA was isolated from Trizol using chloroform extraction, isopropanol and ethanol precipitation. DNase digestion Cleanup along with DNase I digestion was then performed using the RNeasy mini kit (Qiagen). RNA was quantified using nanodrop.

#### Quantification of *maco-1* expression by qPCR

cDNA was made using the Superscript III First-strand system for RT-PCR (Invitrogen). The generated cDNA was used as input for qPCR reactions using the Power SYBR Master Mix and ran on ViiA Real Time PCR System. The primers used are as follows: *maco-1* (Forward: GTGTCACGACAATTGCC; Reverse: CACATAGGTAGTGGCGAG)^4^ and *act-1* (Forward: GGCCCAATCCAAGAGAGGTATC; Reverse: CAACACGAAGCTCATTGTAGAAGG)^53^.

#### Statistical analysis

For choice assays and reversal assays, each dot represents an individual assay plate or individual worm, respectively. Comparisons between two groups used unpaired, two-tailed Student’s *t* tests; comparisons across more than two genotypes used one-way ANOVA with Tukey’s multiple comparisons test; comparisons across genotype × training condition used two-way ANOVA with Tukey’s multiple comparisons test. Experiments were repeated on separate days with independently raised populations to confirm reproducibility. Prism 9 (or later) software was used for all statistical analyses.

## Notes

### Competing Interest Statement

The authors have declared no competing interest.

