## Supplemental Figures for "Glial cells gate small RNA transport from germline to brain during transgenerational epigenetic inheritance of behavior"

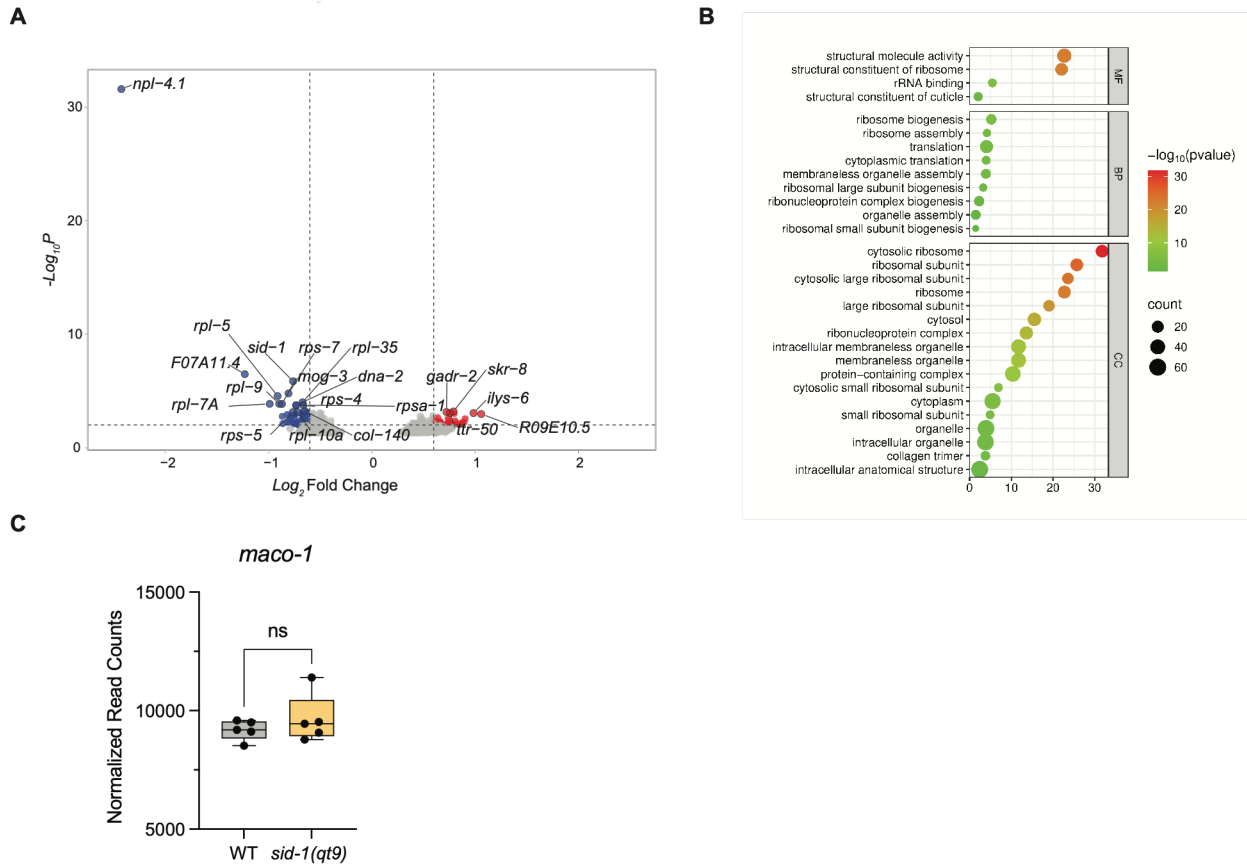

**Figure S1. Whole worm transcriptome analysis reveals numerous differentially expressed genes between WT and *sid-1* animals** **(A)** Volcano plot depicting differential gene expression in wild-type (N2) and *sid-1*(qt9) animals. Genes that are significantly differentially expressed are highlighted in red and blue. **(B)** Wild-type (N2) vs *sid-1*(qt9) RNA-seq GO terms. **(C)** *maco-1* is not downregulated in *sid-1*(qt9) mutants. Boxplot showing the expression levels of *maco-1* in wild-type and *sid-1*(qt9) mutants. Each dot represents a biological replicated, $n = 5$. ns (not significant) $p = 0.3937$ t-test

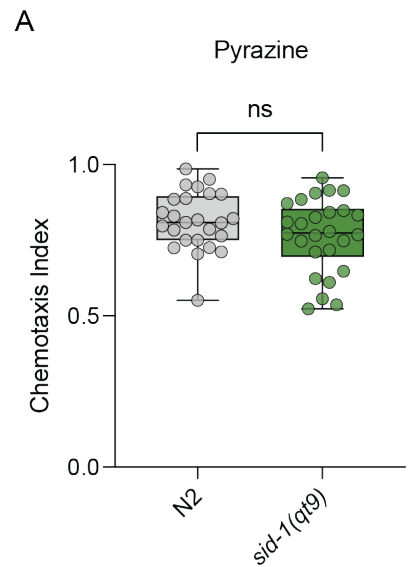

**Figure S2. *sid-1* animals are not defective in pyrazine chemotaxis (A)** Wild-type (N2) vs. *sid-1(qt9)* shows no difference in chemotaxis to pyrazine. Each dot represents a plate with $\geq 3$ 20 worms. $n = 3$ replicates. Center line represents median; box range denotes the 25<sup>th</sup> to 75<sup>th</sup> percentile; whiskers represent minimum to maximum.

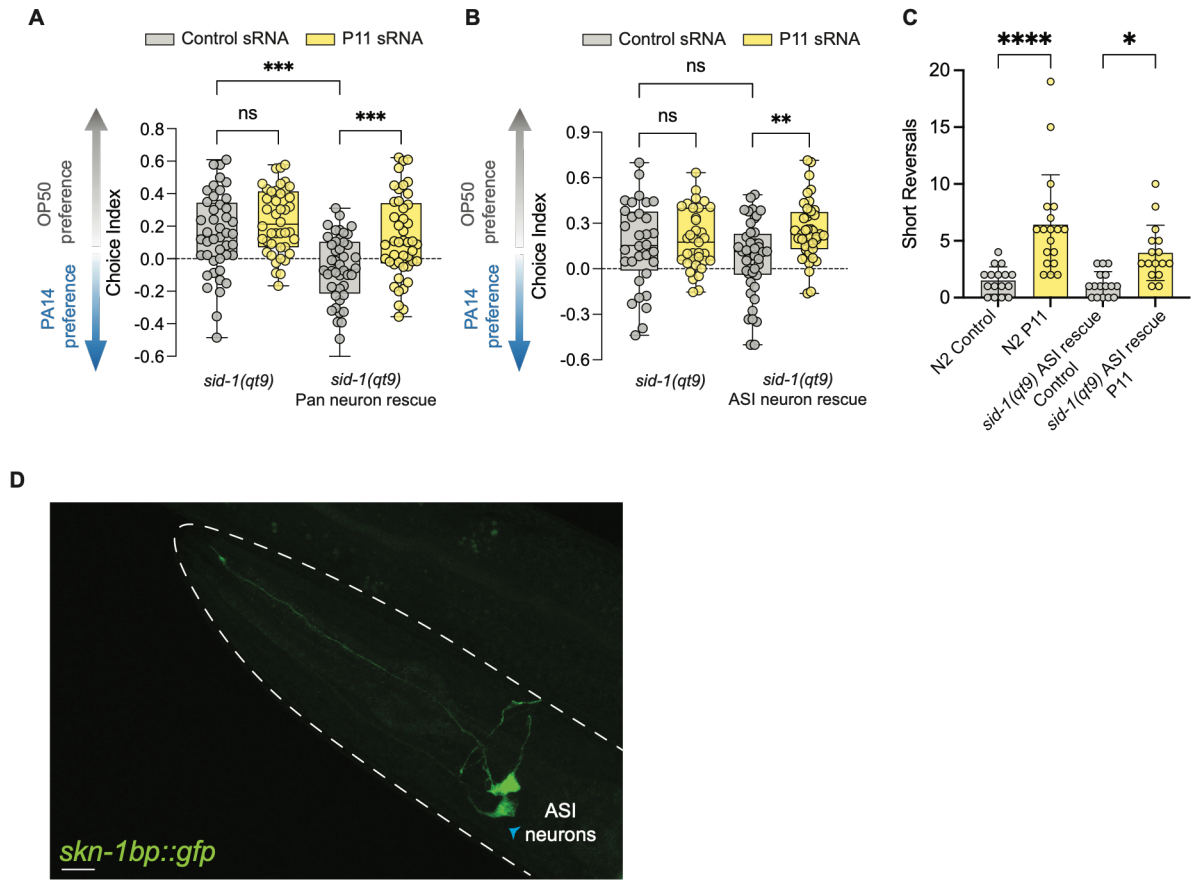

**Fig. S3. Investigating the role of neuronal expression of *sid-1* on learned pathogen avoidances and expression levels of *skn-1b*** (A) *sid-1(qt9)* mutants with *sid-1* rescued in neurons restores naïve attraction to PA14 and P11-mediated learned avoidance. (B) *sid-1(qt9)* mutants with *sid-1* rescued in only in ASI neurons showed restored learning with P11 training. (C) *sid-1(qt9)* mutants with *sid-1* rescued in only in ASI neurons increased short reversals with P11 training. n = 3 biological replicates. \*\*\*\*p<0.0001, \*p<0.05 Two-way ANOVA with Tukey's multiple comparison test. (D) *Pskn-1b::GFP* is expressed in the ASI. Scale bar represents 10μm.

For choice assays each dot represents an assay plate of 31-200 worms, average = 84 worms/plate/genotype/training condition; n ≥ 3 biological replicates. Boxplots: center line represents median; box range denotes the 25<sup>th</sup> and 75<sup>th</sup> percentile; whiskers represent minimum to maximum. Two-way ANOVA with Tukey's multiple comparison test (A-B). \*\*\*p<0.001, \*p<0.05, ns not significant.

**Data S1.**

List of *C. elegans* strains used and generated for this study. Methods used for generation can also be found in this table.

**Data S2.**

Source data file for all figures.
